# Neutralization of Mu and C.1.2 SARS-CoV-2 Variants by Vaccine-elicited Antibodies in Individuals With and Without Previous History of Infection

**DOI:** 10.1101/2021.10.19.463727

**Authors:** Takuya Tada, Hao Zhou, Belinda M. Dcosta, Marie I. Samanovic, Amber Cornelius, Ramin S. Herati, Mark J. Mulligan, Nathaniel R. Landau

## Abstract

Recently identified SARS-CoV-2 variants Mu and C.1.2 have mutations in the receptor binding domain and N- and C-terminal domains that might confer resistance to natural and vaccine-elicited antibody. Analysis with pseudotyped lentiviruses showed that viruses with the Mu and C.1.2 spike proteins were partially resistant to neutralization by antibodies in convalescent sera and those elicited by mRNA and adenoviral vector-based vaccine-elicited antibodies. Virus with the C.1.2 variant spike, which is heavily mutated, was more neutralization-resistant than that of any of variants of concern. The resistance of the C.1.2 spike was caused by a combination of the RBD mutations N501Y, Y449H and E484K and the NTD mutations. Although Mu and C.1.2 were partially resistant to neutralizing antibody, neutralizing titers elicited by mRNA vaccination remained above what is found in convalescent sera and thus are likely to remain protective against severe disease. The neutralizing titers of sera from infection-experienced BNT162b2-vaccinated individuals, those with a history of previous SARS-CoV-2 infection, were as much as 15-fold higher than those of vaccinated individuals without previous infection and effectively neutralized all of the variants. The findings demonstrate that individuals can raise a broadly neutralizing humoral response by generating a polyclonal response to multiple spike protein epitopes that should protect against current and future variants.

## Introduction

SARS-CoV-2 isolates have been classified by the World Health Organization (WHO) as variants of concern (VOC; Alpha (B.1.1.7), Beta (B.1.351), Gamma (B.1.1.248) and Delta (B.1.617.2) and variants of interest (VOI) that include Lambda (C.37)) and newly classified Mu (B.1.621)^1^. In addition, a yet unclassified C.1.2 variant was identified in South Africa^2^ that appears to be increasing in prevalence and spreading to neighboring countries and a variant termed Delta+N501S was identified in Japan, currently at low frequency. Mu^3^ and C.1.2^2,4^ have mutations in the receptor binding domain (RBD) of the spike protein that could contribute to increased transmissibility and cause resistance to neutralization by convalescent sera and vaccine-elicited and therapeutic monoclonal antibodies.

In this study, we measured the infectivity of viruses with the Mu, C.1.2 and Delta+N501S spike proteins and determined their susceptibility to neutralization by convalescent and vaccine-elicited antibodies, both in unexperienced and experienced individuals. We also tested their neutralization by therapeutic monoclonal antibodies. Viruses with the variant spikes were partially resistant to neutralization. The C.1.2 variant, which is highly mutated, was the most resistant. Sera from experienced patients vaccinated with BNT162b2 had very high neutralizing titer against all of the variants, providing a strong rationale for the vaccination of previously infected individuals.

## Results

### Prevalence and infectivity of Mu, C.1.2 and Delta+N501S variants

As of October 2021, the prevalence of the Mu was highest in the British Virgin Islands and Colombia where its accounts for 64% and 43% of sequenced cases **(Figure 1A)**. It is present at low frequency in the Central and South America. The virus has also been found in the United States and Europe although frequencies have not yet been accurately determined. C.1.2 is present with a prevalence of 5% in Swaziland, 1% in South Africa and small numbers of cases have been sequenced in as many as 10 other countries **(Figure 1A)**. In addition, a variant of Delta was recently identified in a handful of cases, termed here Delta+N501S, and has not yet been further characterized.

**Figure 1.**
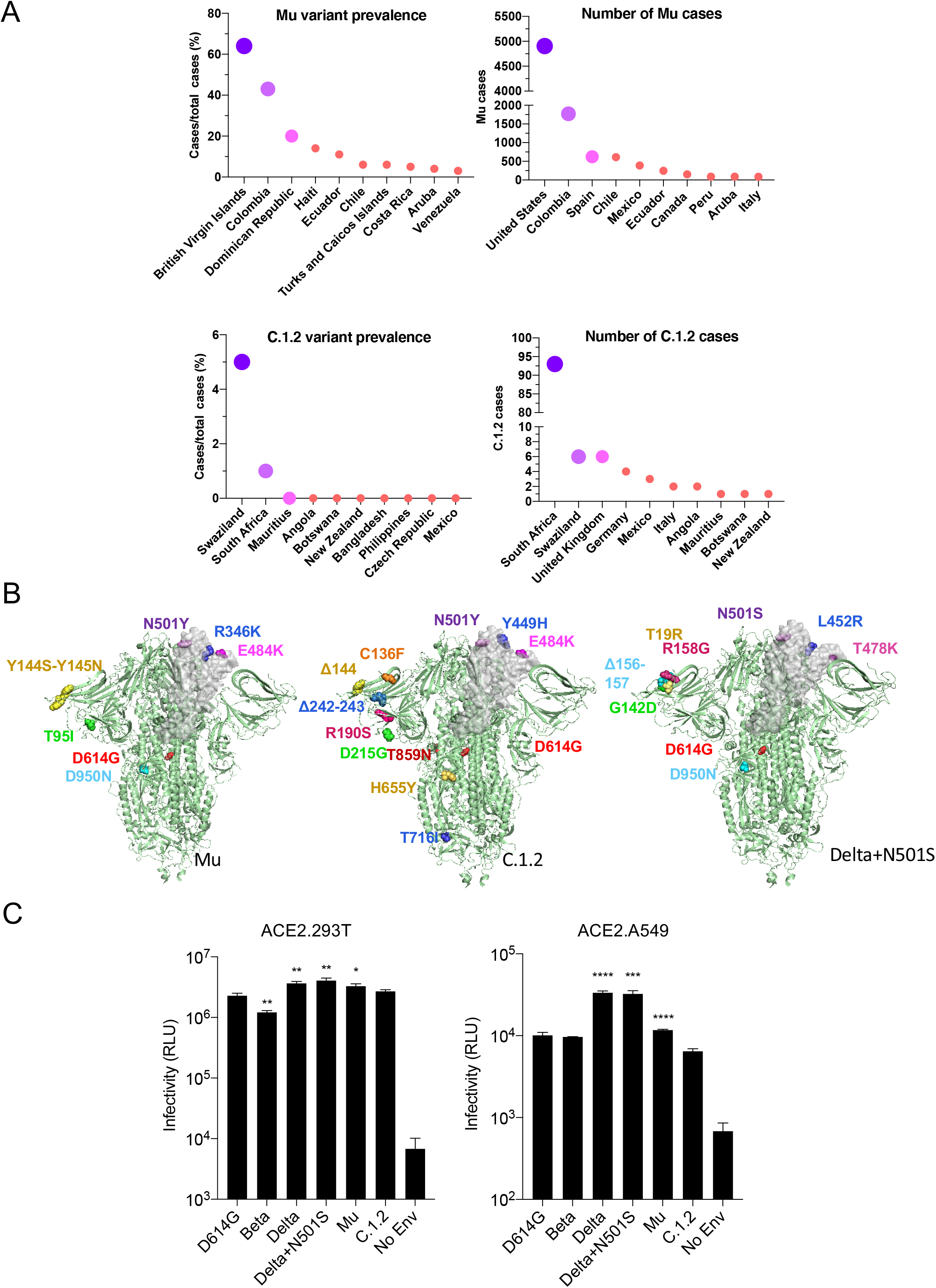
Mu (B.1.621), C.1.2 and Delta+501S variant prevalence and spike protein mutations. (A) The global prevalence of Mu and C.1.2 variants is shown for countries with the highest prevalence or cases (extracted from https://outbreak.info/). (B) Mutations in Mu, C.1.2 and Delta+N501S variant spikes are shown on the three-dimensional spike protein structure. A single RBD in each is shown in gray (side view). The PDB file of spike protein (7BNM)^15^ was downloaded from the Protein Data Bank. 3D view of protein was obtained using PyMOL. (C) The infectivity of Beta, Delta, Delta+N501S, Mu, C.1.2 variant spikes pseudotyped lentiviruses on ACE2.293T and ACE2.A549 cells is shown. The viruses were normalized for RT activity and measured in triplicate with error bars that indicate the standard deviation. The experiment was done three times with similar results.

The variants have unique mutations in the RBD and NTD **(Figure 1B and S1A)**. The Mu spike has RBD mutations R346K, E484K and N501Y; C.1.2 has Y449H, E484K and N501Y; and Delta+N501S has L452R, T478K **(Figure 1B and S1A)**. To evaluate the function and sensitivity of the variant spikes to antibody neutralization, we generated lentiviruses pseudotyped with the Mu, C.1.2 and Delta+N501S spike proteins and, in addition, a pseudotype with the C.1.2 RBD mutations (Y449H, E484K, N501Y) and pseudotypes with the individual RBD mutations of each variant spike. The variant spike proteins were similarly expressed and proteolytically processed in transfected cells and were incorporated into lentiviral virions at a level similar to that of the parental D614G spike protein **(Figure S1B)**.

Analysis of the infectivity of viruses with the variant spike proteins on ACE2.293T and ACE2.A549 cells showed a slight decrease for the Beta spike compared to D614G (1.8-fold) on ACE2.293T cells while Delta, Delta+N501S and Mu were slightly increased **(Figure 1C)**. The pattern of infectivity was similar on ACE2.A549 cells, except that the infectivity differences were somewhat great, most likely due to the low level of ACE2 on these cells. Analysis of the individual point mutations **(Figure S1C)** showed the individual Beta and Delta RBD mutations (R346K, Y449H, E484K, N501Y, N501S) did not significantly increase infectivity. A spike protein with the NTD C.1.2 mutations (P9L-C136F-Δ144-190S-D215G-Δ242-243) also had wild-type infectivity while a spike with the C.1.2 RBD mutations had a significant increase in infectivity (1.9-fold). A spike containing the CTD mutations (H655Y, N679K, T716I, T859) of C.1.2 was decreased 15-fold. Similar infectivity ratios were obtained on ACE2.A549 cells.

### Neutralization of variants by convalescent and vaccine-elicited antibodies

To determine the susceptibility of the viruses with the variant spike proteins to antibody neutralization, we analyzed the neutralizing titers of serum antibodies elicited by the BNT162b2 and mRNA-1273 mRNA vaccines and the Ad26.COV2.S adenoviral vector-based vaccine on the variants. The vaccine sera analyzed were collected from individuals at similar time-points post-final injection, (a mean of 90 days for BNT162b2, 80 for mRNA-1273 and 82 for Ad26.COV2.S; **Table S1**) and all participants tested negative for antibodies against the SARS-CoV-2 N protein suggesting no history of SARS-CoV-2 infection **(Table S1)**. Convalescent sera neutralized D614G spike with a mean titer of 334. Neutralization of Beta, Delta, Delta+ and Mu variants showed a modest 4-9-fold decrease in neutralizing titer while C.1.2 was more resistant to neutralization with a 9-fold decrease **(Figure 2A)**. BNT162b2 sera neutralized virus with the D614G spike with a mean titer of 862, a 2.6-fold increase compared to convalescent sera. The neutralizing titers against Beta, Delta and Delta+N501S were decreased 4.8−, 3.4− and 3.4-fold, respectively. Mu and C.1.2 were somewhat more resistant with a 6.8 and 7.9-fold decrease in titer respectively. mRNA-1273 vaccinated sera showed a similar pattern of neutralization with C.1.2 being the most resistant (11.2-fold decreased titer). Neutralizing antibody titers of sera from Ad26.COV2.S-immunized individuals neutralized D614G with an average titer of 245 and showed a similar pattern of variant neutralization. Titers against C.1.2 fell into a range below 50, the minimum detectable by the assay **(Figure 2B).** Presentation of the data grouped by variant shows decreased neutralizing titers against the variants by sera of the Ad26.COV2.S-vaccinated individuals **(Figure 2C)**. Analysis of the spike proteins with individual variant mutations showed that the neutralization resistance of Mu was caused by R346K and E484K while resistance of C.1.2 was caused by E484K, Y449H and the NTD (P9L-C136F-Δ144-190S-D215G-Δ242-243) **(Figure 2D)**.

**Figure 2.**
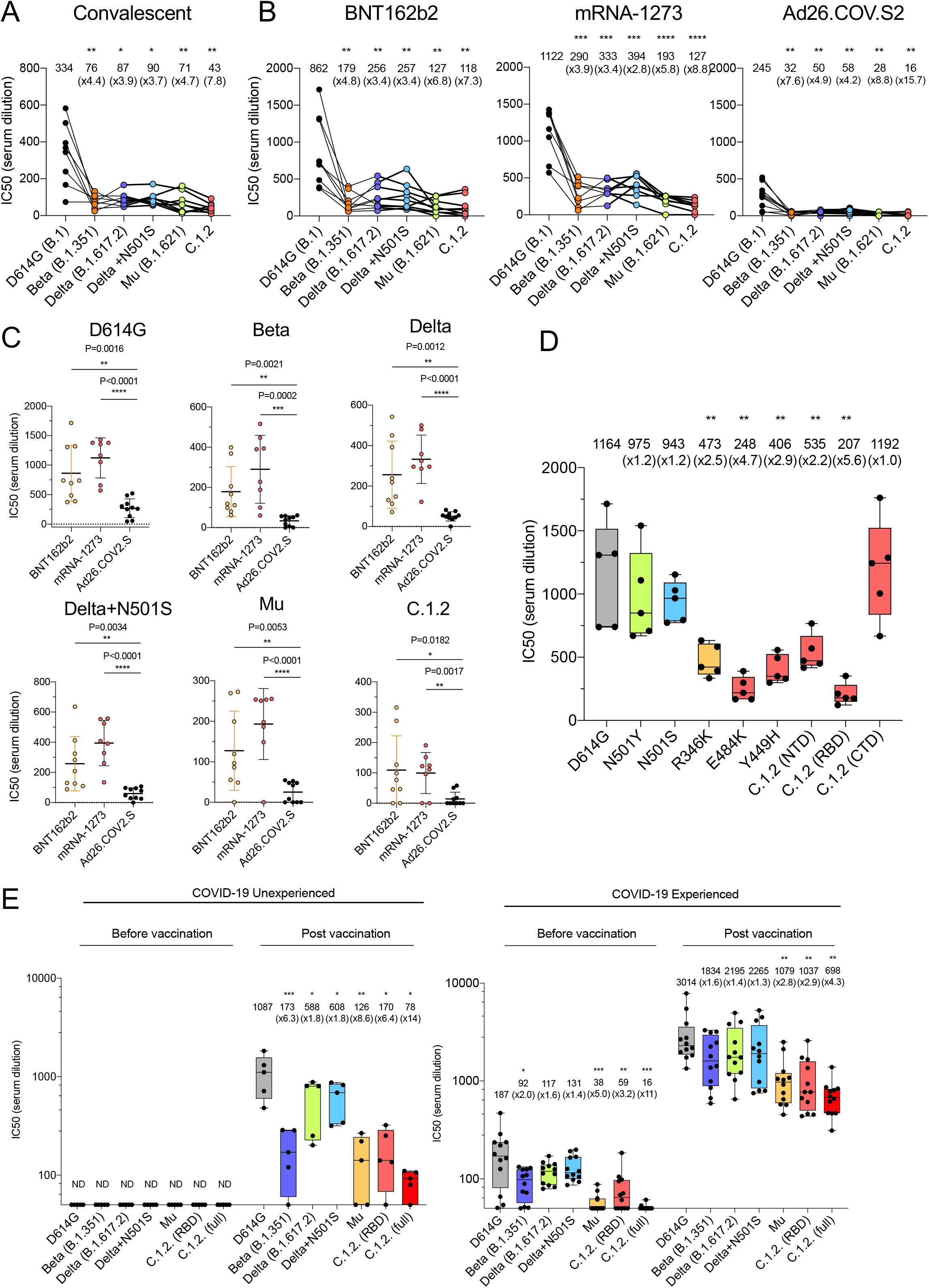
Neutralization of variant spike pseudotyped viruses by convalescent sera, antibodies elicited by RNA and adenoviral vector vaccines. (A) Neutralization of pseudotyped viruses with D614G, Beta, Delta, Delta+N501S, Mu, C.1.2 variant spikes by convalescent serum samples from 8 donors was tested. The serum was collected at 32-57 days after infection. Each dot represents the IC50 for a single donor. Neutralization titers of variants were compared with that of D614G. (B-C) Neutralizing titers of serum samples from BNT162b2 vaccinated individuals (n=9), mRNA-1273 vaccinated donors (n=8), Ad26.COV2.S vaccinated individuals (n=10) was measured. Sera were collected at 90, 80, 82 days on average post-last immunization. IC50 of neutralization of virus from individual donors are shown. Significance was based on two-sided testing. (D) Neutralization titers of viruses with single point mutations by antibodies elicited by BNT162b2. Neutralizing titers of serum samples from BNT162b2 vaccinated individuals (n=5). Aera were collected 7 days post-second immunization. Each dot represents the IC50 for a single donor. (E) Neutralizing titers of serum samples from BNT162b2 vaccinated individuals with (n=5) or without previous SARS-CoV-2 experience (n=12) was measured. The neutralization IC50 of virus from individual donors is shown. The sera were collected 7 days post-second immunization. Significance between variants and D614G was determined by student-t test or Nonparametric ANOVA test. (*P≤0.05, **P≤0.01, ***P≤0.001, ****P≤0.0001). The experiment was done twice with similar results.

Analysis of neutralization by the sera of donors who had a history of COVID-19 pre-BNT162b2 vaccination showed an overall higher neutralizing titer against all of the variants. The neutralizing titer of sera from unexperienced donors against D614G was 1087 on average **(Figure 2E)** with Beta, Delta, Delta+N501S, Mu and C.1.2 having a 1.4-14-fold decrease in titer. In contrast, experienced-vaccinated donor sera were significantly increased in titer against D614G (2.8-fold) and the titers remained high for all of the variants **(Figure 2E)**. After two doses of vaccination, infection-experienced donors had a 10.6-fold increase in neutralizing titer against the Beta variant compared to unexperienced donors. Titers were increased 3.7-fold for Delta and Delta+N501S. Overall, the neutralization titers of sera from experienced donors were 8.5-8.9-fold greater against Mu and C.1.2 variants compared with unexperienced individuals. (**Table S2**)

### Variant spike avidity for ACE2

To measure the ACE2 binding avidity of the variant spikes, we established an ACE2 avidity assay in which the variant spike proteins were expressed in 293T cells and then incubated with a serially diluted soluble ACE2:nanoluciferase fusion protein (sACE2-Nluc) **(Figure 3A)**. Similar cell surface spike protein expression levels on the transfected 293T cells was confirmed by flow cytometry (not shown). The analysis showed increased ACE2 binding affinity of the Beta, Delta, Delta+N501S and Mu spikes (2.2−, 1.7−, 1.4−, 1.7-fold, respectively) as indicated by a decrease in the concentration required to achieve 50% occupancy of the spike protein. In contrast, C.1.2 bound ACE2 with decreased affinity, requiring 1.4-fold higher concentration of ACE2 for 50% binding compared to D614G and 2.4-fold decrease as compared to the high affinity Beta variant spike protein **(Figure 3B)**. Analysis of the point-mutated spike proteins showed that the increased affinity of Beta, Delta, Mu with ACE2 was attributed to N501Y and L452R **(Figure 3B)**, consistent with previous studies^6^. The decreased affinity C.1.2 for ACE2 was due to the combination of the Y449H in the RBD and the mutated NTD. These results were confirmed in a virion binding assay in which pseudotyped virions were incubated with sACE2 and then added to ACE2.293T cells and the amount of bound virions was then measured **(Figure S2)**. In this assay, virions with D614G, Beta, Delta, Delta+N501S, C.1.2 (RBD) and Mu spikes bound similarly to ACE2 while C.1.2 binding was decreased. These findings suggest that the C.1.2 spike protein binds ACE2 with a relatively lower affinity than the other spike protein variants.

**Figure 3.**
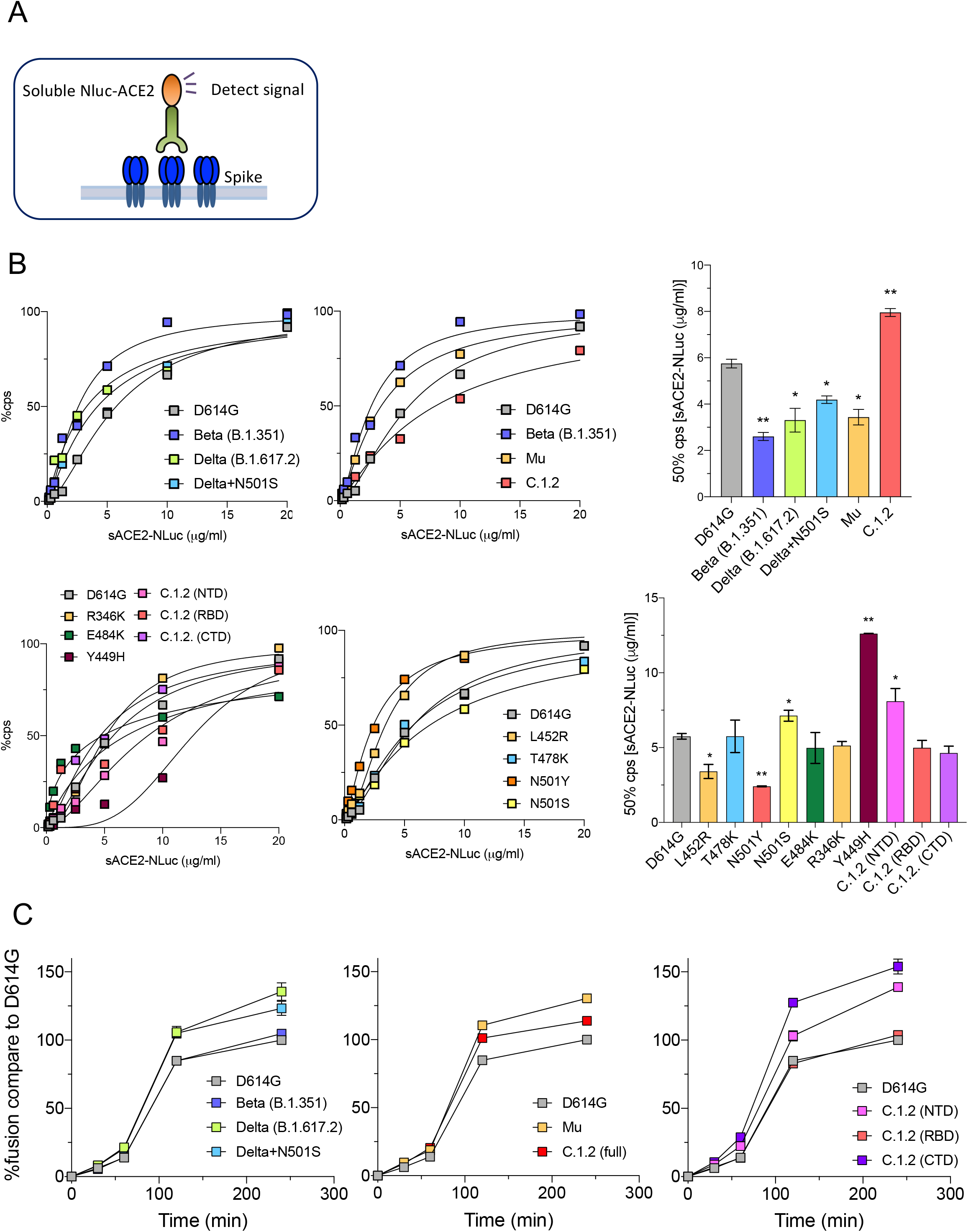
Binding and fusion of variant spikes to ACE2. (A) The diagram shows the principle of the ACE2 avidity assay in which 293T cells transfected with variant spike protein expression vector are incubated with serially diluted sACE2-nluc protein. Following a 30-minute incubation, the unbound fusion protein is removed and the bound protein measured by luciferase assay. (B) ACE2 avidity of the indicated variant spike proteins is shown as curves with maximal binding defined as luciferase activity upon binding of the ACE2.nLuc fusion protein at 50 mg.ml set to 100% (left two panels). The histogram on the right shows 50% of maximal binding. (C) Cell:cell fusion kinetics of the variant spike proteins is shown as measured in an α-complementation assay. The experiment was done twice with similar results.

### SARS-CoV-2 spike protein mediated cell:cell fusion

To test the ability of the variant spike proteins to mediate the fusion reaction upon ACE2 binding, we established an assay for SARS-CoV-2 spike-mediated cell:cell fusion. The assay is based on the alpha-complementation of beta galactosidase strategy that we previously established for the analysis of HIV-1 envelope glycoprotein-mediated fusion^5^. Cells expressing the SARS-CoV-2 spike protein and a peptide were mixed with cells expressing ACE2 and ω fragment **(Figure S3)** and ß-galactosidase activity was measured between 0-5 hours using a luminescent substrate. The results showed that fusion activity could be detected as early as 30 minutes post-mixing and reached a near maximum by 4 hours (>1×10^6^ cps). Analysis of the variant spike proteins with this assay showed that Delta, Delta+N501S, Mu and C.1.2 increased fusion activity compared to D614G **(Figure. 3C)**. Although the fusion activity was same between C.1.2 (RBD) and D614G, C.1.2. (CTD) and C.1.2 (NTD) were higher than D614G, suggesting that the mutations in NTD (P9L-C136F-Δ144-190S-D215G-Δ242-243) and CTD (H655Y, N679K, T716I, T859) affect fusion activity **(Figure 3C)**.

### Therapeutic antibodies neutralize Mu and C.1.2

Regeneron monoclonal antibodies maintained their ability to neutralize Delta, Delta+N501S, Mu and C.1.2. REGN10933 lost titer (50-fold) against virus with the Beta spike **(Figure S4A)**, as previously reported^6–8^ but maintained neutralizing activity against the others while REGN10987 maintained activity against all variants **(Figure S4B)** and the combination of the two mAbs was highly active against all of the variants **(Figure S4C)**.

## Discussion

Virus with the newly described Mu and C.1.2 spike proteins were partially resistant to neutralization by antibodies in convalescent sera and to those elicited by mRNA and adenoviral vector-based vaccine-elicited antibodies. The C.1.2 variant spike, which is heavily mutated, was the most neutralization-resistant of the variant spike proteins tested here and was more resistant than those on which we have previously reported^6^. The resistance of the C.1.2 spike was caused by a combination of the RBD mutations N501Y, Y449H and E484K and the NTD mutations.

Mathematical modeling by Khoury *et al*. predicts that 50% protection from SARS-CoV-2 infection is provided by a titer that is 20% that of the convalescent titer^9^. In this study, mean convalescent titer was 334 (**Table S1**), indicating that 50% protection would correspond to an IC50 of 67. The titer required to protect against severe disease is predicted to be 3% that of the mean titer of convalescent sera, corresponding to a titer of 10 in this study, suggesting that vaccination should remain protective against severe disease resulting from infection with Mu or C.1.2.

Interestingly, the neutralizing titers of sera from infection-experienced BNT162b2-vaccinated individuals, those with a history of previous SARS-CoV-2 infection, were on average 6.4-fold higher than those of vaccinated individuals without previous infection and effectively neutralized all of the variants. The findings demonstrate that individuals can raise a broadly neutralizing humoral response^10,11^, presumably by generating a polyclonal response to multiple spike protein epitopes, that will protect against current and most likely, future variants. In addition, Regeneron therapeutic monoclonal antibodies retained their ability to neutralize Mu and C.1.2 variants.

An unexplained finding in our analysis regards the effect of the T716I mutation in the C.1.2 spike. The mutation in the C.1.2 spike caused an 18-fold decrease in infectivity which was alleviated when the mutation was taken out and replaced with a threonine at position 716 (**Figure S1D**). The mutation is also present in the Alpha variant spike protein where it similarly caused a marked decrease in the infectivity^6^. To ensure that the mutation did not affect the findings of our study, the C.1.2 spike protein used here contained all of the mutations except T716I. The reason for decreased infectivity of the fully mutated C.1.2 spike is unclear. It may be a result of producing the spike protein in 293T cells perhaps caused by the proximity of T716 to a potential glycosylation site at position 717. The T716I mutation had no effect on expression of the protein in transfected cells, packaging into virions or ACE2 avidity. The mutation also had no effects on antibody neutralization profile (data not shown). Thus, the T716I mutation did not influence the results but this effect should be considered in studies with pseudotypes using spike proteins with this mutation.

It is interesting that the C.1.2 spike protein has a decreased affinity for ACE2 compared to that of the other variant spikes. Its RBD has the N501Y mutation that confers increased ACE2 affinity^12,13^ but this is counteracted by the novel Y449H mutation that decreases ACE2 affinity. The decreased ACE2 affinity is unexpected as it is thought the virus is mutating to increase ACE2 affinity and thereby increase transmissibility. This would suggest that Y449H was selected as an antibody escape mutation, a finding consistent with the increased resistance to antibody neutralization indicated in our data. The decrease in ACE2 affinity of the C.1.2 spike protein may cause a decrease in transmissibility, and thereby limit spread of the virus, despite its relative resistance to antibody neutralization. It suggests that as the virus is selected to escape the humoral immune response, it becomes less fit, unable to evolve a highly transmissible, neutralization resistant variant.

The Mu variant does not appear to present any additional concerns over Delta with which it is nearly identical. The C.1.2 variant is currently at low prevalence and has a restricted geographic distribution but given the large number of NTD mutations coupled with RBD and CTD mutations and relative neutralization resistance, the spread of the variant should be closely monitored. The high titers of antibody against all of the variants in experienced patients is encouraging because it demonstrates the ability of individuals to mount a broadly neutralizing antibody response that will may be impervious to current and future variants. By extrapolation, the finding suggests that vaccine booster immunization might result in a similarly broad antibody response.

**Table 2.** IC5O of sera from unexperienced and experienced individuals before and after vaccination with BNTI62b2 against viruses with variant spike proteins. Age, Sex and Comorbidities are shown.

| Covid Unexperienced BNT162b2 |  |  |  |  |  |  |  |  |  |  |  |
| --- | --- | --- | --- | --- | --- | --- | --- | --- | --- | --- | --- |
| Note: Covid experienced samples had no detectable levels of antibodies against variants before vaccination |  |  |  |  |  |  |  |  |  |  |  |
| donor | Days post 2<sup>nd</sup> dose | Age | Sex | Comorbidities | IC<sub>50</sub>(serum dilution) (post vaccination) |  |  |  |  |  |  |
|  |  |  |  |  | D614G | Beta | Delta | Delta +N501S | Mu | C.1.2 (RBD) | C.1.2 (full) |
| 1 | 8 | 62 | Male | none | 1824 | 285 | 882 | 871 | 141 | 136 | 80 |
| 2 | 8 | 34 | Male | none | 1307 | 171 | 802 | 825 | 267 | 322 | 0 |
| 3 | 8 | 40 | Female | none | 1109 | 287 | 804 | 687 | 221 | 141 | 93 |
| 4 | 8 | 39 | Female | none | 713 | 120 | 251 | 317 | 0 | 251 | 110 |
| 5 | 9 | 39 | Male | none | 481 | 0 | 201 | 341 | 0 | 0 | 107 |
| Mean (SD) | 8.2 (0.4) | 43 (11) |  |  | 1087 (524) | 173 (121) | 588 (332) | 608 (264) | 126 (123) | 170 (123) | 78 (45) |

|  |  |  |  |  | IC<sub>50</sub>(serum dilution) |  |  |  |  |  |  |  |  |  |  |  |  |  |
| --- | --- | --- | --- | --- | --- | --- | --- | --- | --- | --- | --- | --- | --- | --- | --- | --- | --- | --- |
| donor | Days post 2<sup>nd</sup> dose | Age | Sex | Comorbidities | D614G |  | Beta |  | Delta |  | Delta +N501S |  | Mu |  | C.1.2 |  | C.1.2 (full) |  |
|  |  |  |  |  | Before | After | Before | After | Before | After | Before | After | Before | After | Before | After | Before | After |
| 1 | 10 | 34 | M | none | 66 | 5477 | 107 | 2462 | 129 | 1957 | 165 | 2157 | 31 | 755 | 69 | 1660 | 29 | 838 |
| 2 | 11 | 46 | M | none | 163 | 7844 | 51 | 3183 | 171 | 4964 | 192 | 4157 | 17 | 2118 | 71 | 2585 | 3 | 660 |
| 3 | 7 | 60 | M | none | 287 | 3852 | 129 | 3133 | 125 | 3823 | 120 | 5268 | 76 | 2501 | 184 | 1731 | 53 | 807 |
| 4 | 9 | 42 | F | none | 470 | 2179 | 40 | 853 | 90 | 652 | 86 | 752 | 62 | 570 | 0 | 939 | 0 | 863 |
| 5 | 12 | 43 | M | none | 237 | 1829 | 51 | 1421 | 79 | 1539 | 111 | 979 | 0 | 1247 | 61 | 771 | 0 | 311 |
| 6 | 8 | 37 | F | none | 236 | 2760 | 89 | 3299 | 112 | 4007 | 105 | 4488 | 63 | 1029 | 94 | 668 | 61 | 696 |
| 7 | 10 | 54 | F | Hypertension, History of obesity | 156 | 2101 | 125 | 2246 | 148 | 1178 | 168 | 794 | 23 | 1051 | 105 | 461 | 12 | 468 |
| 8 | 7 | 34 | F | none | 220 | 1739 | 72 | 1368 | 141 | 1218 | 199 | 1592 | 11 | 1080 | 98 | 454 | 0 | 468 |
| 9 | 7 | 32 | F | Anemia, Asthma, GERD | 54 | 1970 | 74 | 667 | 86 | 1025 | 107 | 797 | 88 | 453 | 0 | 437 | 22 | 471 |
| 10 | 11 | 24 | F | none | 179 | 2412 | 109 | 1793 | 114 | 2485 | 93 | 2027 | 20 | 577 | 15 | 771 | 13 | 785 |
| 11 | 10 | 49 | F | Hypertension, Rhinosinusitis | 126 | 1343 | 133 | 591 | 135 | 1742 | 127 | 2395 | 35 | 651 | 9 | 610 | 5 | 618 |
| 12 | 6 | 44 | F | Hereditary Hemorrhagic Telangiectasias, Eosinophilic Esophagitis | 51 | 2664 | 125 | 993 | 82 | 1754 | 98 | 1783 | 28 | 921 | 0 | 1357 | 0 | 1388 |
| Mean (SD) | 9 (2) | 42 (11) |  |  | 187 (118) | 3014 (1885) | 92 (34) | 1834 (1008) | 118 (29) | 2195 (1359) | 131 (39) | 2266 (1554) | 38 (28) | 1079 (629) | 59 (57) | 1037 (665) | 17 (21) | 698 (279) |

## Acknowledgements

The work was funded by grants from the NIH to N.R.L. (DA046100, AI122390 and AI120898) and to M.J.M. (UM1AI148574).

## Author contributions

T.T., H.Z. and N.R.L. designed the experiments. T.T., H.Z. and B.M.D. carried out the experiments and analyzed data. T.T., H.Z., B.M.D. and N.R.L. wrote the manuscript. M.I.S., A.C., R.H. and M.J.M supervised specimen selection and the collection of clinical information, did the ELISAs and provided reagents and key insights. All authors provided critical comments on manuscript.

---

**Supplementary Figure 1.**
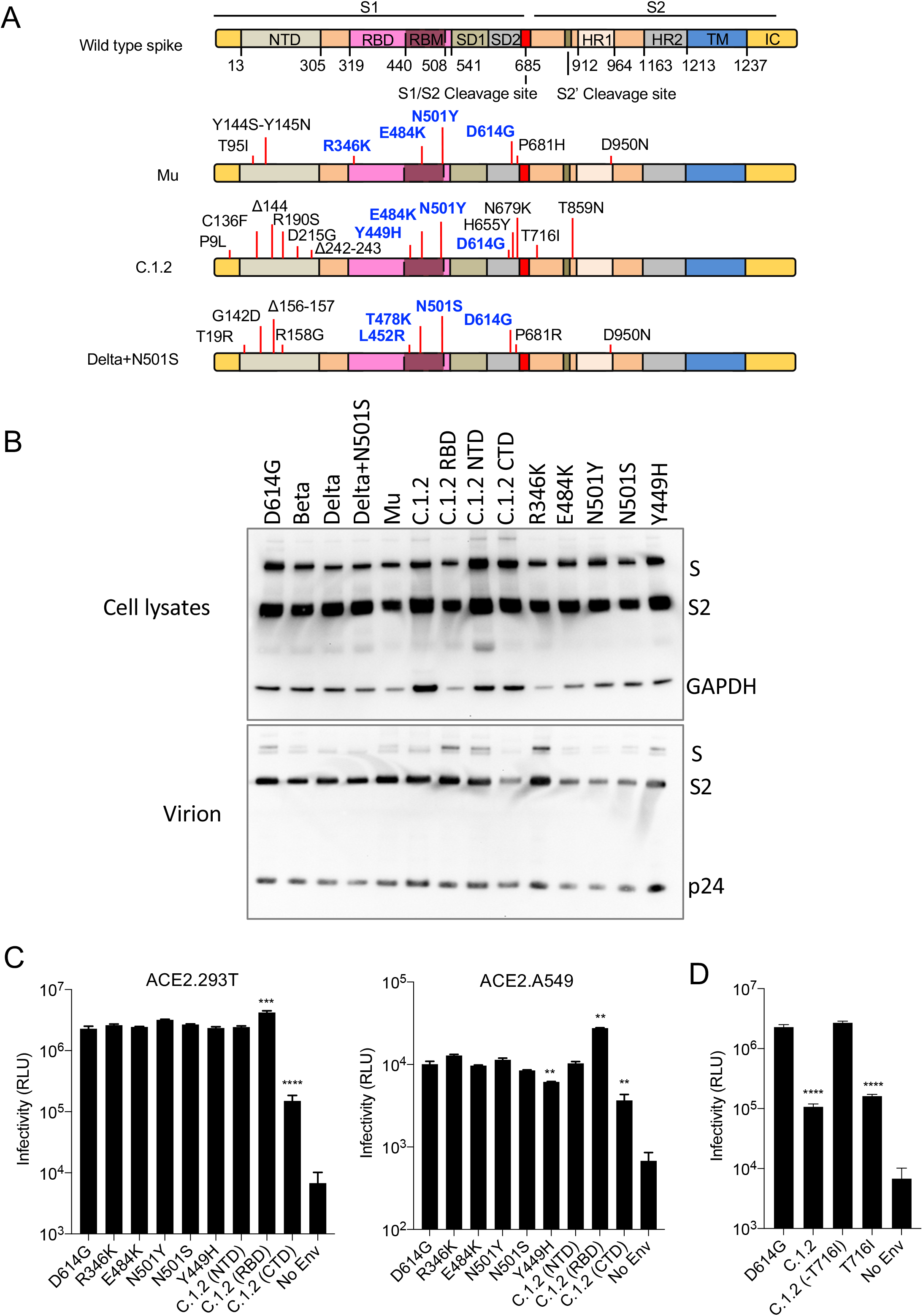
Infectivity of variant spike viruses and spike expression levels. (A) The domain structure of the SARS-CoV-2 spikes of Mu (B.1.621), C.1.2 and Delta+N501S is diagrammed. NTD, N-terminal domain; RBD, receptor-binding domain; RBM, receptor-binding motif; SD1 subdomain 1; SD2, subdomain 2; HR1, heptad repeat 1; HR2, heptad repeat 2; TM, transmembrane region; IC, intracellular domain. (B) Immunoblot analysis of the variant spike proteins in transfected 293T cells. Pseudotyped viruses were produced by transfection of 293T cells. Two days post-transfection, virions were analyzed on an immunoblot probed with anti-spike antibody and anti-HIV-1 p24. The cell lysates were probed with anti-spike antibody and anti-GAPDH antibodies as a loading control. (C) Infectivity of virus pseudotyped by Beta, Delta, Delta+N501S, Mu, C.1.2 variant individual spikes pseudotyped lentivirus in ACE2.293T and ACE2.A549 cells. (D) Infectivity of virus pseudotyped by C.1.2 (full), C.1.2. (-T716I) variant spikes and T716I variant individual spike pseudotyped lentivirus in ACE2.293T cells. The experiment was done twice with similar results.

**Supplementary Figure 2.**
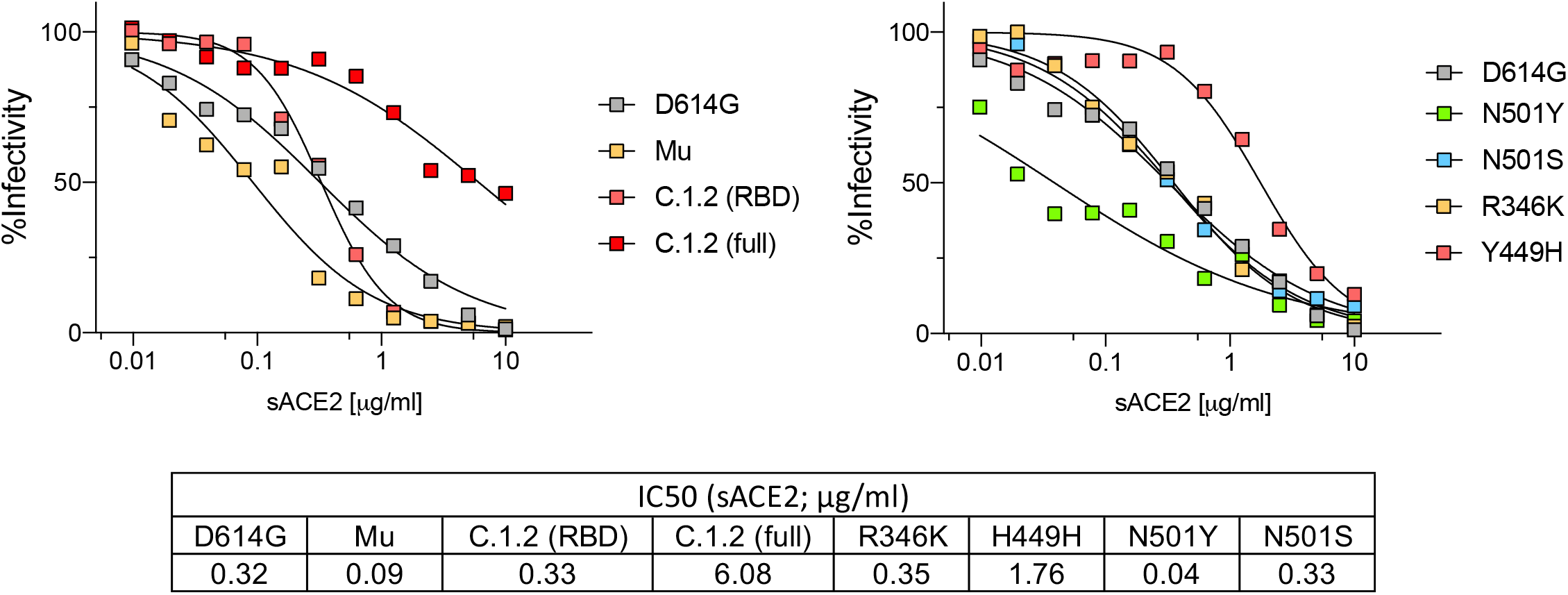
Neutralization of spike protein variants by sACE2. Serially diluted sACE2 was mixed with variant pseudotypes (D614G, Mu, C.1.2) (left) and single point mutated virus (N501Y, N501S, R346K, Y449H) (right) for 30 minutes. The mixture was added on ACE2.293T cells. After 2 days of infection, luciferase activity was measured. The experiment was done three times with similar results.

**Supplementary Figure 3.**
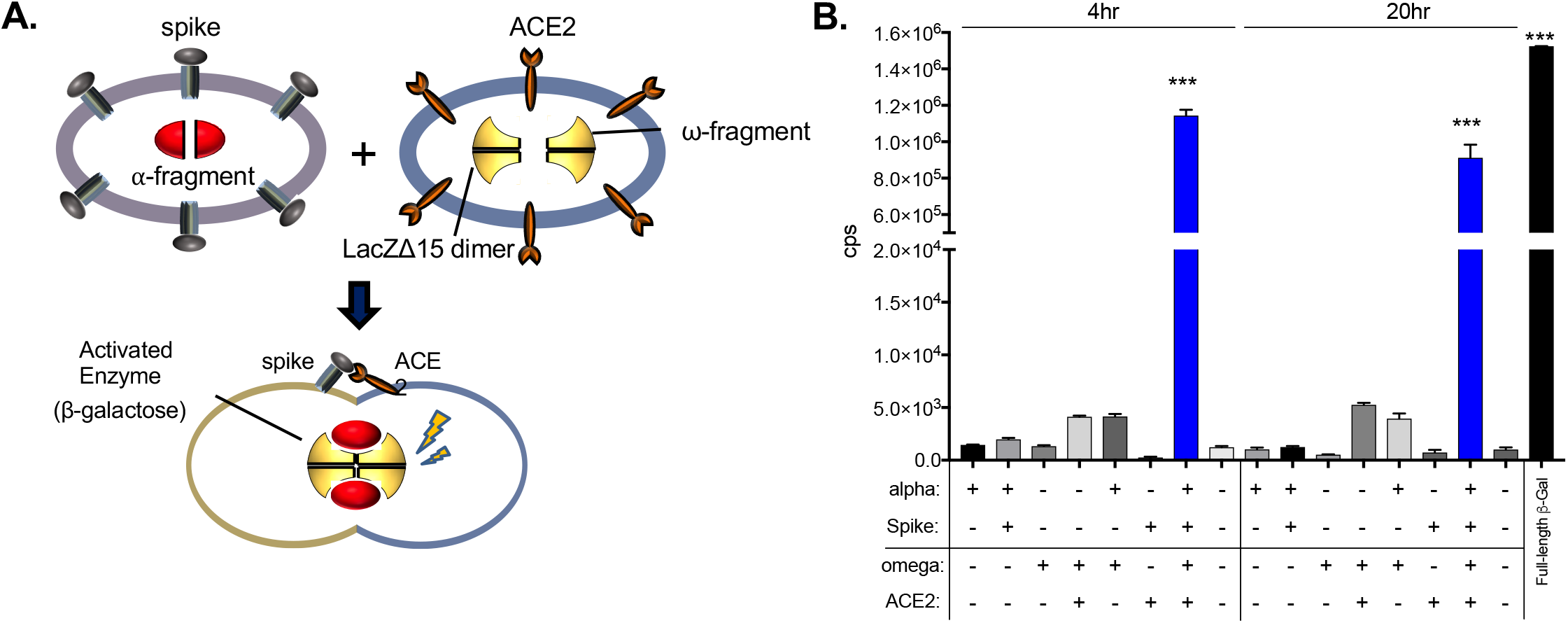
α-Complementation cell:cell fusion assay. (A) The assay is based on the enzymatically inactive β-galactosidase α peptide (red) and C-terminal ω fragment (yellow). The fragments are expressed separately in transfected cells. Upon mixing of cells separately expressing a spike protein and ACE2, enzymatically active β-galactosidase tetramers are formed. Effector 293T cells that express alpha-N85 and SARS-CoV-2-Δ19 spike were incubated with target 293T cells that express ω and ACE2. ß-galactosidase activity was measured after 4 and 20 hours of incubation. (B) 293T cells expressing alpha and spike were mixed with target cells. After 4 and 20 hours of incubation, ß-galactosidase activity was measured. The data are displayed as the mean ± SD and significance as calculated in the student-t test.

**Supplementary Figure 4.**
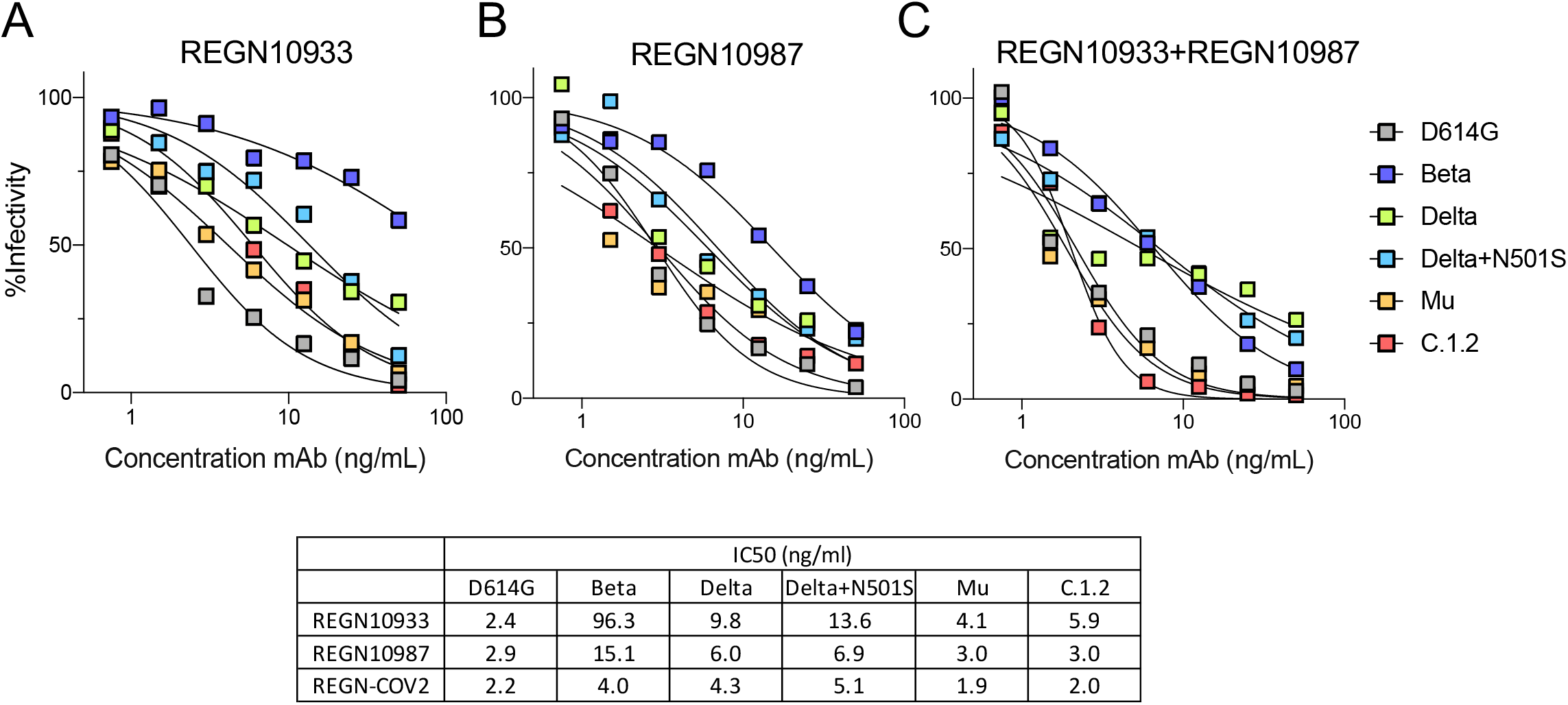
Neutralization of spike protein variants by monoclonal antibodies REGN10933 and REGN10987. (A-C) Neutralization of D614G, Beta, Delta, Delta+N501S, Mu, C.1.2 variant spikes by REGN10933 and REGN10987. Neutralization of viruses by REGN10933 (A), REGN10987 (B), and 1:1 mixture of REGN10933 and REGN10987 (C) was measured. The table shows the calculated IC50 for each curve. The experiment was done three times with similar results.

## Methods

### Plasmids

Plasmids used in the production of lentiviral pseudotyped virus have been previously described^14^. Mutations were introduced into pcCOV2.Δ19.D614G by overlap extension PCR and confirmed by DNA sequencing.

### Human sera and monoclonal antibodies

Convalescent sera were collected 32-57 days post-symptom onset. BNT162b2-vaccinated sera were collected 90 days (mean) post-second immunization and mRNA-1273-vaccinated sera were collected 80 (mean) days post-second immunization. Ad26.COV2.S-vaccinated sera were collected 82 days (mean) post-immunization. COVID-19 experienced serum samples were collected 7 days post-second immunization with BNT162b2. Participants reported experiencing COVID symptoms were confirmed COVID-19-experienced by direct PCR or antibody testing. The clinical study was conducted at the NYU Vaccine Center with participant’s written consent under IRB-approved protocols (18-02035 and 18-02037). sACE2 was generated as previously described^14^.

### SARS-CoV-2 spike lentiviral pseudotypes

Lentivirus pseudotyped by variant SARS-CoV-2 spikes were produced as previously reported^14^. Viruses were concentrated by ultracentrifugation and normalized for reverse transcriptase (RT) activity. Sera and monoclonal antibodies were serially diluted and then incubated with pseudotyped virus (approximately 2.5 × 10^7^ cps) for 30 minutes at room temperature and then added to target cells. Luciferase activity was measured 2 days postinfection.

### Binding assay

293T cells were transfected with mutated spike variant expression vectors using lipofectamine 2000 and seeded in a 96-well plate at 1 × 10^4^ / well. Serially diluted sACE2 protein fused with nano-Luciferase was added to the cells. Following incubation for 30 minutes at 37°C, the unbound proteins were washed and luciferase activity was measured using Nano-Glo substrate (Nanolight) in an Envision 2103 microplate luminometer (PerkinElmer).

### Neutralization assay by soluble ACE2

Briefly, pseudotyped virus was incubated with serially diluted recombinant soluble ACE2 protein for 1 hour at room temperature and subsequently added to 1 × 10^4^ ACE2.293T cells. After 2 days, the cell medium was removed and 50 μl Nano-Glo luciferase substrate (Nanolight) was added. The luminescence signal was read in an Envision 2103 microplate luminometer.

### α-complementation assay

293T (4 × 10^6^) were cotransfected with pcΔ19S and 5 μg pSCTZ-alpha N85 or variable amounts of pLenti.ACE2-HA and 5 μg of pSCTZ-omega by lipofection with lipofectamine 2000 (Invitrogen). After 24 hours, the medium was changed and the following day, the transfected cells were collected with PBS/5 mM EDTA. Spike protein+alpha-N85 transfected cells (2 × 10^5^) were incubated with patient serum or ACE2 microbody for 30 min at room temperature and then mixed with an equal number of ACE2+omega transfected cells in a volume of 100 μl in a 96-well culture plate. After 4 hours, β-galactosidase activity was measured using the Galacto-Light Plus β-Galactosidase Reporter Gene Assay System (Thermo Fisher). The cells were lysed in 100 μl Tropix Lysis Buffer for 10 min at room temperature and then 20 μl of the lysate was mixed with 70 μl Galacto-Plus substrate diluted 1:100 in Tropix Galacto Reaction Buffer Diluent. After incubation for 30 min at room temperature, 100 μl of Tropix Accelerator II was added and the luminescence was read in an Envision 2103 microplate luminometer (PerkinElmer). For β-Galactosidase detection with FDG, the cells were lysed in 100 μl of buffer containing 10 mM Tris pH 8.0, 150 mM NaCl and 0.1% triton X-100. After 5 min, 10 μl of the lysate was mixed with 100 μl of Tropix Galacto Reaction Buffer Diluent and 10 μl of 2 mM FDG (Thermo Fisher) in 50% DMSO. The reactions were incubated for 30 min in the dark after which luminescence was visualized by illumination with 365 nm UV light or on an iBright gel documentation instrument (Thermo Fisher).

### Immunoblot analysis

Proteins were analyzed on immunoblots probed with mouse anti-spike monoclonal antibody (1A9) (GeneTex), anti-p24 monoclonal antibody (AG3.0) and anti-GAPDH monoclonal antibody (Life Technologies) followed by goat anti-mouse HRP-conjugated secondary antibody (Sigma).

### Statistical Analysis

All experiments were performed in technical duplicates or triplicates and the data were analyzed using GraphPad Prism 8. Statistical significance was determined by the two-tailed unpaired t-test or Nonparametric ANOVA test. Significance was based on two-sided testing and attributed to p< 0.05. Confidence intervals are shown as the mean ± SD or SEM (*P≤0.05, **P≤0.01, ***P≤0.001, ****P≤0.0001).

---

**Table S1.**
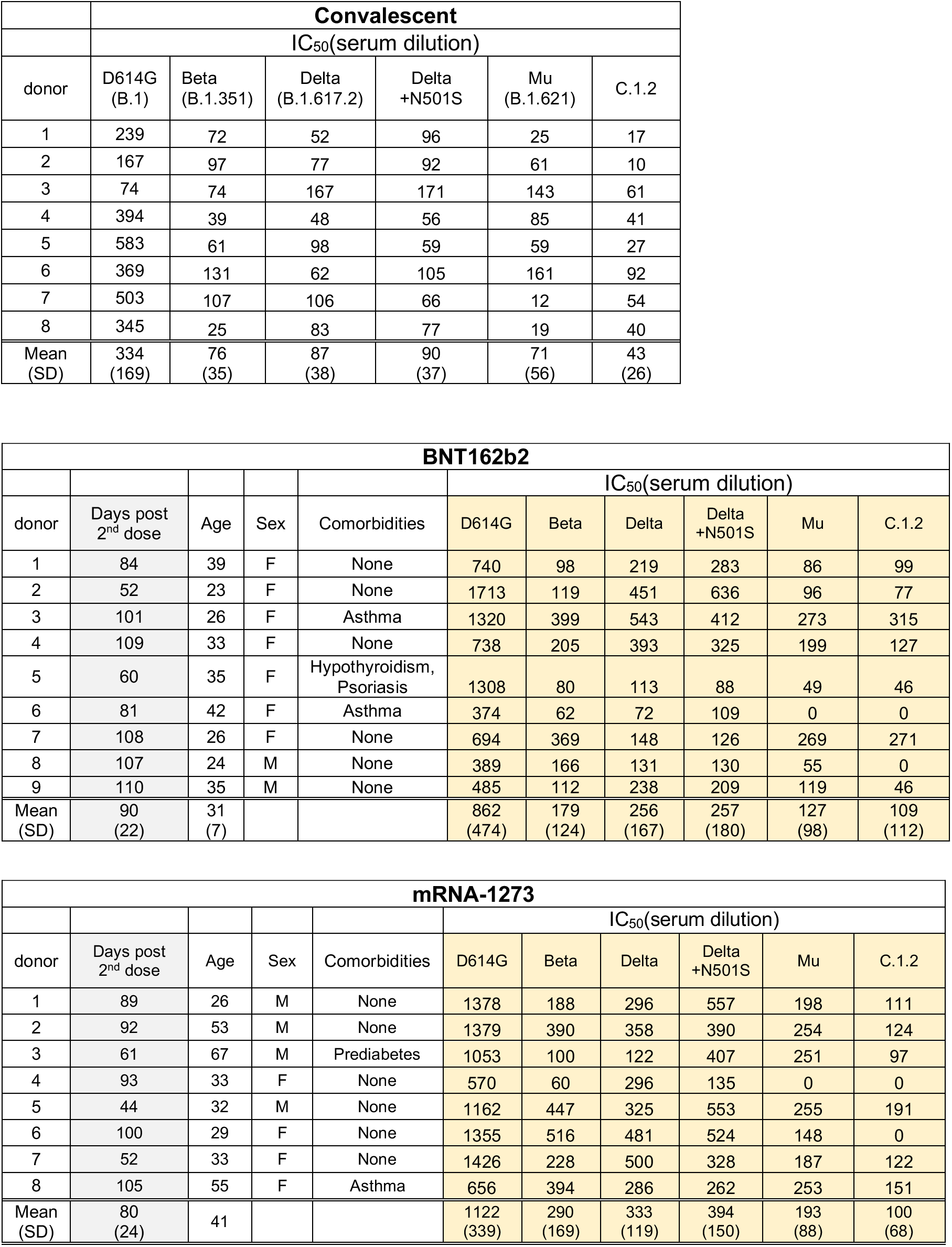
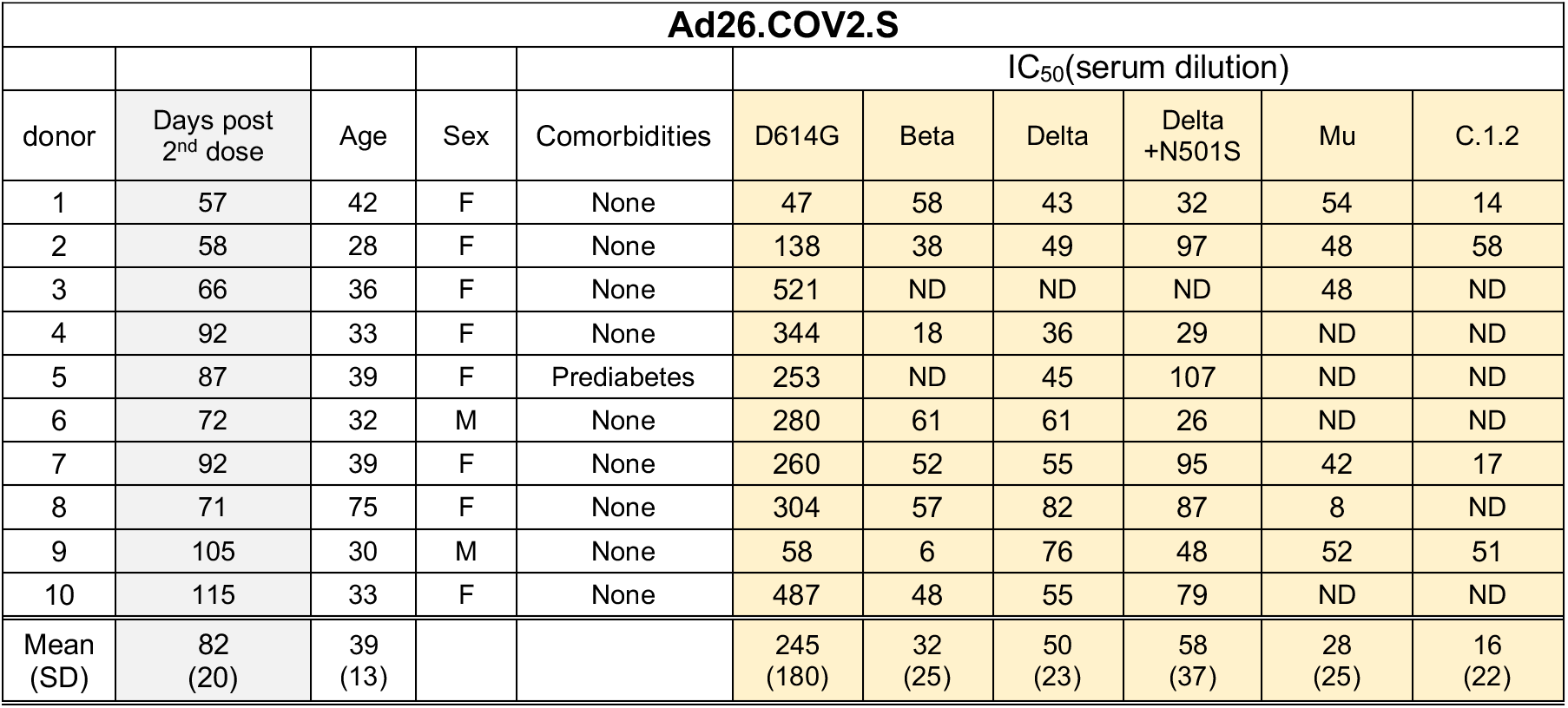
IC5O of Convalescent, BNTI62b2, mRNA-1273 and Ad26.COV.S elicited antibodies against viruses with variant spike proteins. Age, Sex and Comorbidities are shown.

